# Antisense Oligonucleotide Targeting Hepatic Serum Amyloid A Limits the Progression of Angiotensin II-Induced Abdominal Aortic Aneurysm Formation

**DOI:** 10.1101/2023.08.22.554377

**Authors:** Preetha Shridas, Ailing Ji, Andrea C Trumbauer, Victoria P Noffsinger, Luke W Meredith, Frederick C de Beer, Adam E Mullick, Nancy R Webb, Dennis G Karounos, Lisa R Tannock

## Abstract

**Background and aim:** Obesity increases the risk for abdominal aortic aneurysms (AAA) in humans and enhances angiotensin II (AngII)-induced AAA formation in C57BL/6 mice. We reported that deficiency of SAA significantly reduces AngII-induced inflammation and AAA in both hyperlipidemic apoE-deficient and obese C57BL/6 mice. The aim of this study is to investigate whether SAA plays a role in the progression of early AAA in obese C57BL/6 mice.

**Methods:** Male C57BL/6J mice were fed a high-fat diet (60% kcal as fat) throughout the study. After 4 months of diet, the mice were infused with AngII until the end of the study. Mice with at least a 25% increase in the luminal diameter of the abdominal aorta after 4 weeks of AngII infusion were stratified into 2 groups. The first group received a control antisense oligonucleotide (Ctr ASO), and the second group received ASO that suppresses SAA (SAA-ASO) until the end of the study.

**Results:** Plasma SAA levels were significantly reduced by the SAA ASO treatment. While mice that received the control ASO had continued aortic dilation throughout the AngII infusion periods, the mice that received SAA-ASO had a significant reduction in the progression of aortic dilation, which was associated with significant reductions in matrix metalloprotease activities, decreased macrophage infiltration and decreased elastin breaks in the abdominal aortas.

**Conclusions:** We demonstrate for the first time that suppression of SAA protects obese C57BL/6 mice from the progression of AngII-induced AAA. Suppression of SAA may be a therapeutic approach to limit AAA progression.

## Introduction

Abdominal aortic aneurysm (AAA) is a degenerative vascular disorder associated with sudden death due to aortic rupture. It affects up to 8% of men over 65 in the United States, with an annual mortality of more than 15,000 ^1^. AAA becomes a more serious public health issue as the elderly population increases.

Surgical therapies have shown no benefit in the treatment of small aortic aneurysms (<5 cm), as the risk of rupture is comparable to the risks of surgical intervention. This asymptomatic interval of “watchful waiting” provides an opportunity for medical intervention to reduce AAA expansion and, hence, the risk of rupture. Unfortunately, despite multiple clinical trials, no therapy has proven effective in blunting AAA progression. Thus, there is an urgent need for better therapies ^2^. An in-depth exploration of the mechanisms underlying AAA occurrence and inhibition is needed to support the development of potent treatments. AAA involves the dilation and thinning of the arterial wall in the abdominal segment of the aorta ^3^. Macrophage infiltration, inflammatory cytokine release, and matrix metalloproteinase (MMP) activation are factors mediating the medial injury and adventitial inflammation, ultimately leading to the aneurysmal dilation of the aorta in AAA ^4, 5^.

Serum Amyloid A (SAA) is an acute-phase protein whose plasma concentrations can increase > 1000-fold and exceed 1 mg/ml during acute inflammation. However, SAA is also persistently elevated in chronic inflammatory conditions such as obesity ^6, 7^, diabetes ^6, 8^, and atherosclerosis ^9^. In humans, acute-phase SAAs comprise SAA1 and SAA2 (SAA1.1 and SAA2.1 in mice), and SAA3 is an additional acute-phase SAA in mice (SAA3 is a pseudogene in humans) ^10^. Mouse SAA proteins are highly homologous to their human counterparts ^11^. SAA is considered to be predominantly produced by the liver ^12^.

Chronic infusion of Angiotensin II (AngII) to hyperlipidemic or obese male mice produces progressive luminal dilation of the abdominal aorta and pathology closely resembling human AAA ^13, 14^. AngII infusion induces systemic SAA in mice. Our group reported the key finding that deficiency of SAA in hyperlipidemic apolipoprotein E-deficient (apoE^-/-^) mice significantly reduces AngII-induced AAA ^15^. Recently, we also reported that SAA exacerbates AAA development in obese mice infused with AngII ^16^. In the present study, we investigated whether antisense-oligonucleotide (ASO)-mediated inhibition of SAA protects mice from the progression of an early AAA.

## Materials and Methods

### Animal Treatments

All procedures involving animals were approved by the Institutional Animal Care and Use Committees at the University of Kentucky and/or the Lexington Veterans Affairs Medical Center. All experiments were performed in male mice as they develop aneurysms at a significantly higher incidence compared to female mice, similar to the clinical observation that AAAs predominantly affect men ^17^. C57BL/6J mice (8-week-old) were obtained from The Jackson Laboratory and fed an obesogenic diet (60% kcal from fat; D12492, Research Diets) for 28 weeks, as indicated in Fig.1. AngII (1000 ng/kg/min; 4006473, BACHEM) was continuously infused via Alzet osmotic minipumps (model 2004; Durect Corporation) for the last 12 weeks of diet with a change in pumps every 4 weeks. GalNAc-antisense oligonucleotide (ASO, Ionis Pharmaceuticals) solutions in PBS were *i.p.* injected into mice (2 injections in the first week and 1 injection per week in subsequent weeks for 8 weeks at 5 mg/kg/injection). Body composition was measured by NMR spectroscopy (Echo MRI; Echo Medical Systems, Houston, TX, USA). Blood pressure was measured on 5 consecutive days in conscious mice using a non-invasive tail-cuff method (CODA 8; Kent Scientific Corp, Torrington, CT) as described ^18^. The timeline for all the procedures is depicted in Fig.1.

**Figure 1.**
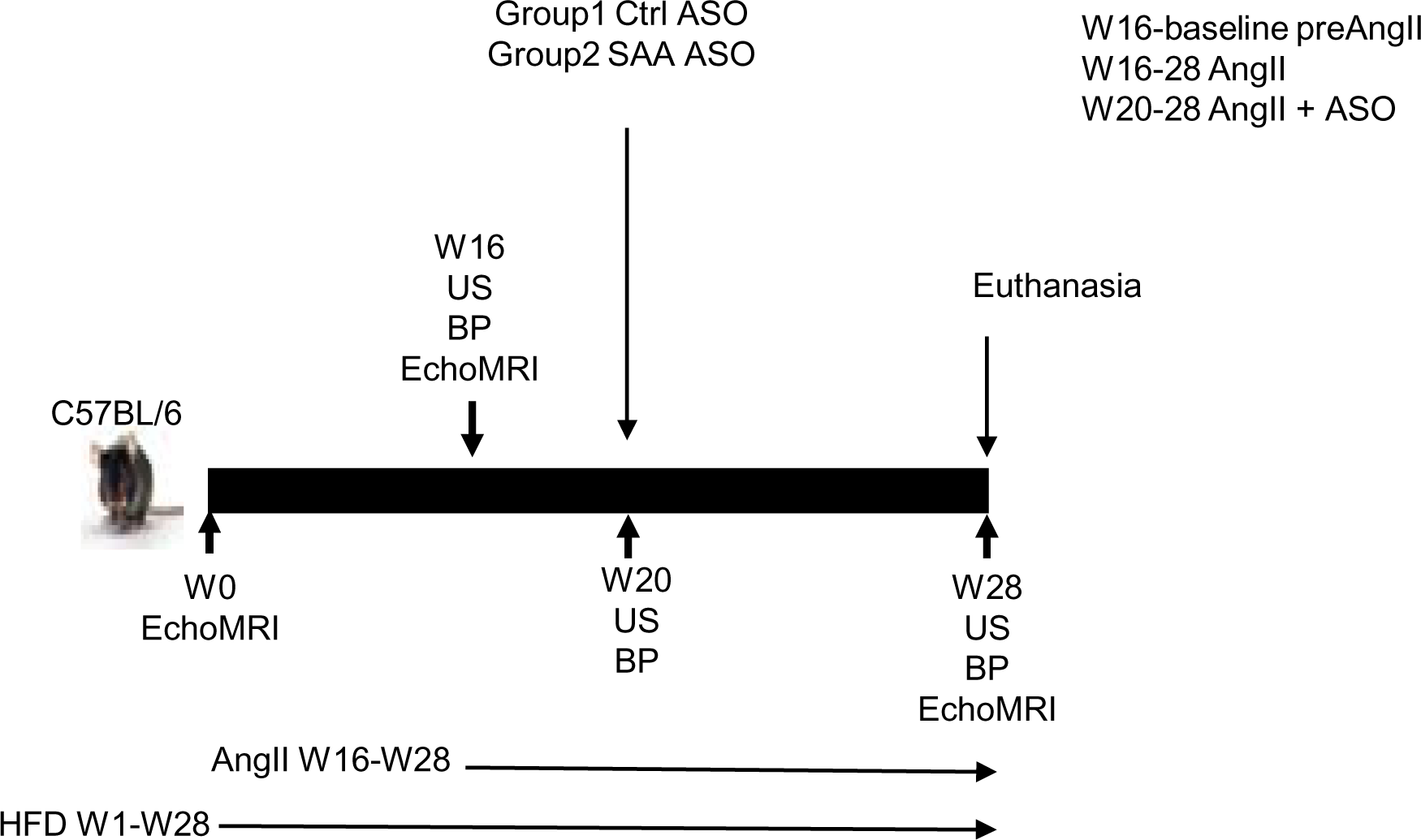
Study design and timeline (weeks). The experiment was carried out in two waves, 8-week-old male C57BL/6J mice (n=30/experiment) were used for each study wave. US: ultrasound analysis for determining the aortic diameter, BP: blood pressure.

### Quantification of AAA

Ultrasound measurements were performed in mice before and after AngII infusions as indicated in Fig.1. Abdominal aortas were visualized in mice anesthetized with 2% v/v Isofluorane with medical grade oxygen (Butler Schein) using high-frequency ultrasound (US) (Vevo 660; VisualSonics) as described previously ^19^. Briefly, anesthetized mice were restrained in a supine position for ultrasonography, and short axis scans of abdominal aortas from the level of the left renal arterial branch moving vertically to the suprarenal region were obtained. Cine loops of 100 frames were acquired throughout the renal region of the abdominal aortas. Maximal luminal diameter and area measurements were determined by two different observers blinded to groups.

### Plasma measurements

Plasma SAA (SAA1.1 and SAA2.1 isoforms) concentrations were determined using a mouse SAA ELISA kit (cat no TP 802M, Tridelta Development Ltd). Plasma cholesterol concentrations were measured using enzymatic kits (Wako Chemicals).

### Immunohistochemistry (IHC)

Immunofluorescence staining was performed as described previously ^15^. Abdominal aortae were frozen in Optimal Cutting Temperature compound (4583, OCT; Tissue-Tek), and 8 μm thick sections down the length of the aorta were mounted on glass slides. All tissue sections were subjected to identical processing at the same time to allow for direct comparison. Sections were fixed in 4% paraformaldehyde for 30 minutes and treated with 0.1% Triton X-100 in PBS for 15 minutes. After blocking in 1% BSA/PBS at room temperature for 2 hr, slides were incubated overnight at 4°C with a combination of rabbit anti-mouse SAA (1:200; ab199030, Abcam) and rat anti-mouse CD68 (1:200; ab53444, Abcam). After washes with PBS, SAA was detected using Alexa Fluor 568–labeled goat anti-rabbit IgG (1:200; A11011, Thermo Fischer Scientific) and CD68 was detected using Alexa Fluor 488–labeled goat anti-rat IgG (1:200; A11006, Thermo Fischer Scientific). Slides were mounted using a fluorescence-protecting medium containing DAPI (Vectashield; Vector Laboratories). Images were captured by ZEISS Axioscan 7 (ZEISS Microscopy) and quantified using Nikon NIS-elements software.

### Elastin staining

For elastin staining, OCT-embedded abdominal aorta sections were fixed in 10% formalin and treated according to the manufacturer’s instructions (Elastic Stain Kit, AB150667, Abcam). Images were captured on a Nikon ECLIPSE 80i microscope with the aid of NIS-Elements BR 4.00.08 software.

### In situ zymography

In situ zymography was performed as described earlier ^15^. Briefly, OCT-embedded abdominal aorta sections, adjacent to those used for IHC, were incubated with 20 μg/ml DQ gelatin fluorescein conjugate for 2 h at 37°C according to kit instructions (EnzChek Gelatinase/Collagenase Assay Kit, E-12055, Molecular Probes, Inc). The fluorescence generated by hydrolysis of the added substrate was captured by ZEISS Axioscan 7 (ZEISS Microscopy) and quantified using Nikon NIS-elements software. The general MMP inhibitor 1,10-phenanthroline, 20 mmol/L, was used to define non-specific fluorescence.

### RNA isolation and quantitative RT-PCR

Total RNA was isolated from mouse liver tissues according to the manufacturer’s instructions (RNeasy® Mini Kit, 74106, Qiagen). RNA samples were incubated with DNase I (79254, Qiagen) for 15 min at RT prior to reverse transcription. Liver tissue RNA (0.5 μg) was reverse transcribed into cDNA using the Reverse Transcription System (4368814, Applied Biosystems). After 4-fold dilution, 5 µl was used as a template for real-time RT-PCR. Amplification was done for 40 cycles using Power SYBR Green PCR master Mix Kit (4367659, Applied Biosystems). Quantification of mRNA was performed using the ΔΔCT method and normalized to GAPDH. Primer sequences will be provided on request.

### Statistical analyses

Results are expressed as the mean ± SEM, as indicated in the figure legends. Statistics were calculated with GraphPad’s Prism 9 software (GraphPad Software, Inc.). For experiments containing two groups, results were analyzed by Student’s *t*-test. To compare multiple groups, Kruskal-Wallis one-way ANOVA on Ranks followed by Dunn’s method was used. For multiple groups with two factors, two-way ANOVA with the Sidak multiple comparisons test was used. P < 0.05 was considered statistically significant. Fisher’s exact test was applied to the comparisons of AAA incidence. *P* values less than 0.05 were considered statistically significant and denoted in figures with a single asterisk (∗); *P* values less than 0.01 and 0.001 were denoted with two (∗∗) and three asterisks (∗∗∗), respectively.

## Results

To investigate whether SAA contributes to the progression of an early AAA, we used an Anti-Sense Oligonucleotide (ASO)-mediated gene suppression strategy. The experimental design for this study is illustrated in Fig.1. Male C57BL/6J mice were fed a high-fat diet (HFD, 60% kcal as fat) throughout the study. After ∼4 months of diet (W16), the mice were infused with angiotensin II (AngII) at 1000ng/kg/min until euthanasia (W28). Ultrasound (US) was performed in all mice before and after 28 days of AngII infusion. Mice that had at least a 25% increase in the luminal diameter compared to the respective baseline luminal diameter of the abdominal aorta were stratified by luminal diameter and then alternatively assigned to one of two treatment groups based on numerical rankings: a control antisense oligonucleotide (Ctr-ASO) or an ASO that suppresses acute phase SAA isoforms SAA1.1 and SAA2.1 (SAA-ASO) as described in Materials and Methods. Mice continued to receive AngII for a further 8 weeks along with either the Ctr-ASO or SAA-ASO. US was repeated at the study end to assess AAA progression. Mice that did not meet the 25% increase in luminal diameter after 4 weeks of AngII compared to the respective baseline diameters were dropped from the study, and the data generated from these mice are excluded from further analysis.

### SAA ASO administration significantly suppressed plasma SAA levels and liver SAA1.1 and SAA2.1 expression

HFD feeding and AngII infusion increases plasma SAA levels in mice^15,16^. As expected, intraperitoneal (*i.p.*) administration of SAA ASO resulted in a significant reduction in plasma SAA (SAA1.1 and SAA2.1) levels (Fig. 2A; 89.2 ± 25.08 µg/ml for the Ctr-ASO group vs. 18.6 ± 0.21 µg/ml for the SAA-ASO group). Consistently, liver SAA1.1 and SAA2.1, but not SAA3, mRNA expression was significantly reduced in mice treated with SAA ASO compared to mice treated with control ASO when analyzed at the end of the study (Fig. 2B and C).

**Figure 2.**
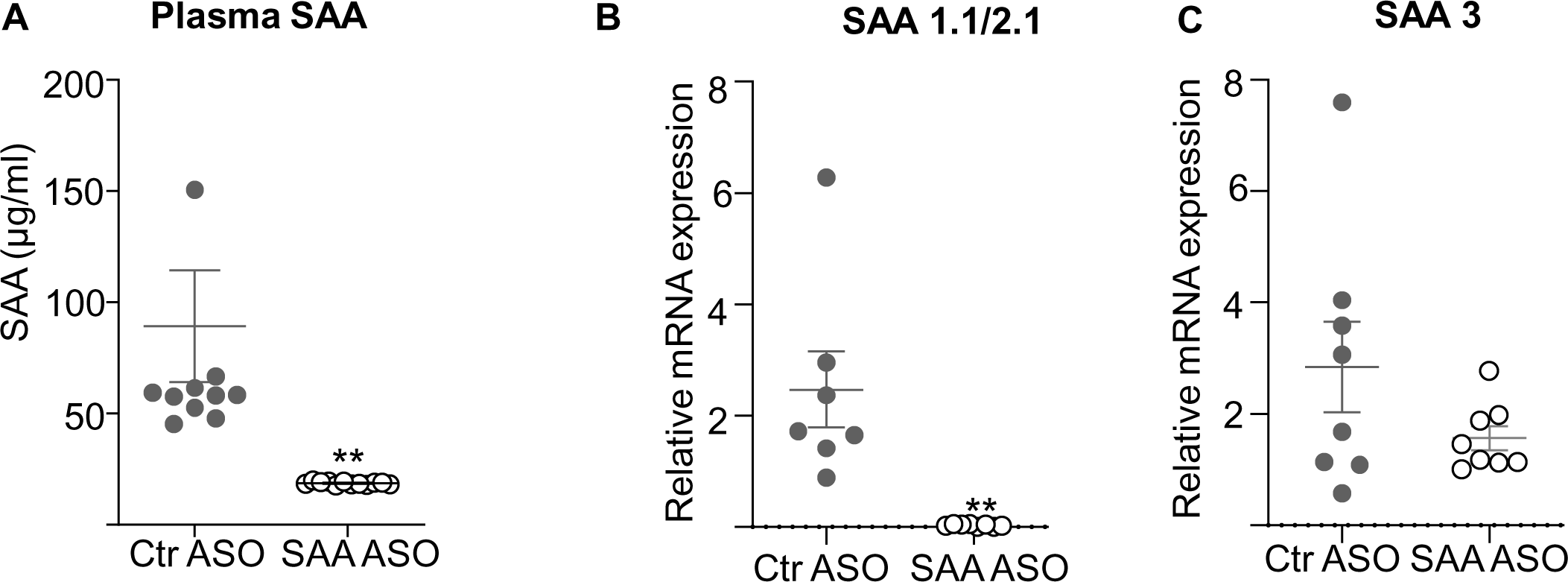
Validation of the effectiveness of ASO for suppressing SAA expression. Experimental mice were treated with antisense oligonucleotides (ASO), either control ASO or SAA ASO (GalNAc SAA-ASO), for the last 8 weeks of the study. (A) Plasma SAA levels at the end of the study were determined by ELISA as described in methods (B) SAA1.1/SAA2.1 and (C) SAA3 mRNA expressions in the livers were determined by qPCR. Data are mean ±SEM from n=6-10 mice/group. **= P ≤ 0.01.

### Suppression of SAA does not change body weight, blood pressure, plasma cholesterol, or lipoprotein profile

The study showed no significant differences in body weights between mice treated with Ctr ASO and SAA ASO during the course of the study (Fig. 3A). As expected, obesogenic diet feeding increased body weights in mice, with an average increase in body fat content from 4.9% to 37.5% before and after 16 weeks of diet, respectively (data not shown). All the mice lost weight following the first pump implantation, as previously observed in the AngII infusion model^15^. The subsequent pump implantations did not result in any significant change in body weights (Fig. 3A). There was no significant difference in body fat content between Ctr ASO and SAA ASO groups when analyzed at week-28 (8 weeks post ASO treatment) (data not shown). Mean arterial blood pressure (MAP) increased significantly following AngII infusion in all mice (week-16 vs. week-20 in Fig.3B). There was a trend for a decrease in MAP following SAA ASO treatment; however, the change was not significant compared to the Ctr ASO group (Fig. 3B, week-28). Plasma cholesterol levels were not significantly different between the Ctr ASO group and SAA ASO group at the end of the study (Fig. 3C). Consistently, there were no significant differences in lipoprotein profiles between the two groups (Fig. 3D). Analysis of systemic proinflammatory markers indicated no detectable levels of TNFα or IL-1β, the levels of IL-6 were not significantly different before (8.56±5.1 pg/ml) and after Ctr ASO (2.7±0.79 pg/ml) or SAA ASO (3.8±1.72 pg/ml) treatments.

**Figure 3.**
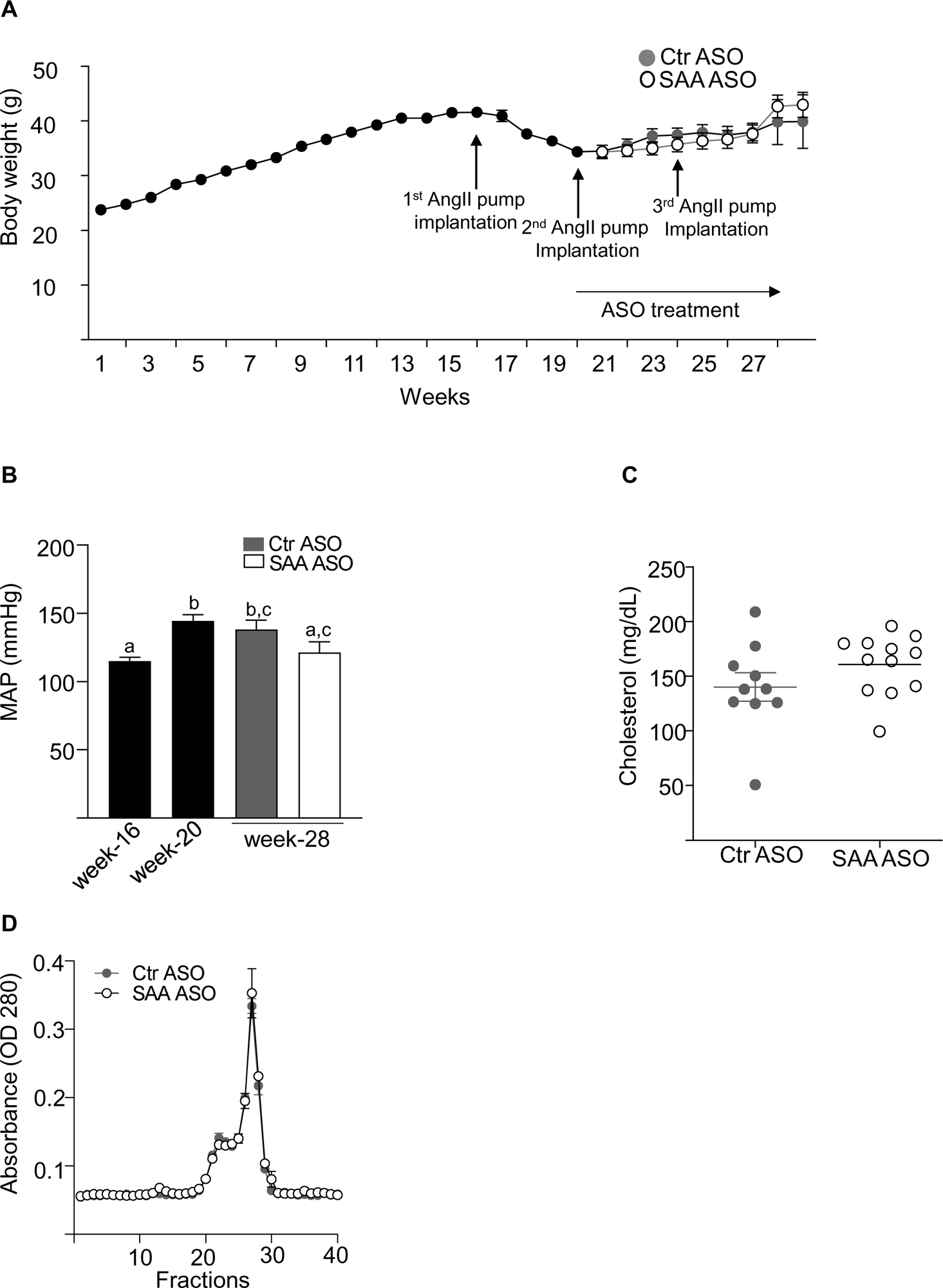
Suppression of SAA did not affect the body weight changes, blood pressure, plasma cholesterol, or lipoprotein profile. (A) body weight changes during the course of the study. The time of AngII infusion and ASO treatment are indicated by arrows. Data are mean ±SEM from n=20 mice/group. (B) Blood pressure was measured on 5 consecutive days in conscious mice using a non-invasive tail-cuff method as described in the methods. Mean arterial pressure (MAP) measurements were obtained prior to osmotic pump implantation (week-16) and 4-(week-20) and 12 (week-28) weeks after pump implantation. Data are mean ±SEM from n=20 mice/group. Groups that are not significantly different (p ≥ 0.05) are indicated with the same letter. (C) Plasma cholesterol concentrations were determined in mice treated with control ASO or SAA ASO for the last 8 weeks of the study, as described in the methods. Data are mean ±SEM from n=10 mice/group. (D) Four plasma samples, chosen from each group’s median range of cholesterol concentration, were pooled for FPLC analysis to determine the lipoprotein profile (pooled from n=4/group).

### Suppression of SAA limits AngII-induced AAA progression

The baseline average aortic luminal diameter for the mice was 1.174±0.026 mm (mean±SEM) prior to AngII infusion. After the first 4 weeks of AngII infusion, the average luminal diameter in all mice was 1.861±0.094 mm (mean±SEM). The average aortic luminal diameter for the mice assigned to receive Ctr ASO (closed square, week-20; Fig. 4A) and SAA ASO (open square, week-20; Fig. 4A) groups after 4 weeks of AngII treatment were 1.885±0.134 and 1.838±0.1356 mm, respectively. Mice that received the control ASO had continued aortic dilation with continued AngII infusion (closed square, week-28; average luminal aortic diameter 2.06±0.12 mm); however, the mice that received the SAA-ASO had a significant reduction in progression of aortic dilation (open square, week 28; average luminal diameter 1.64±0.13 mm), p=0.0015 for interaction between time and group. Representative US images are shown in Fig. 4B. Consistently, the maximal external diameter of the abdominal aorta decreased significantly in mice treated with SAA ASO (1.241±0.04) compared to Ctr ASO (1.879±0.07; Fig. 4C, p<0.0001).

**Figure 4.**
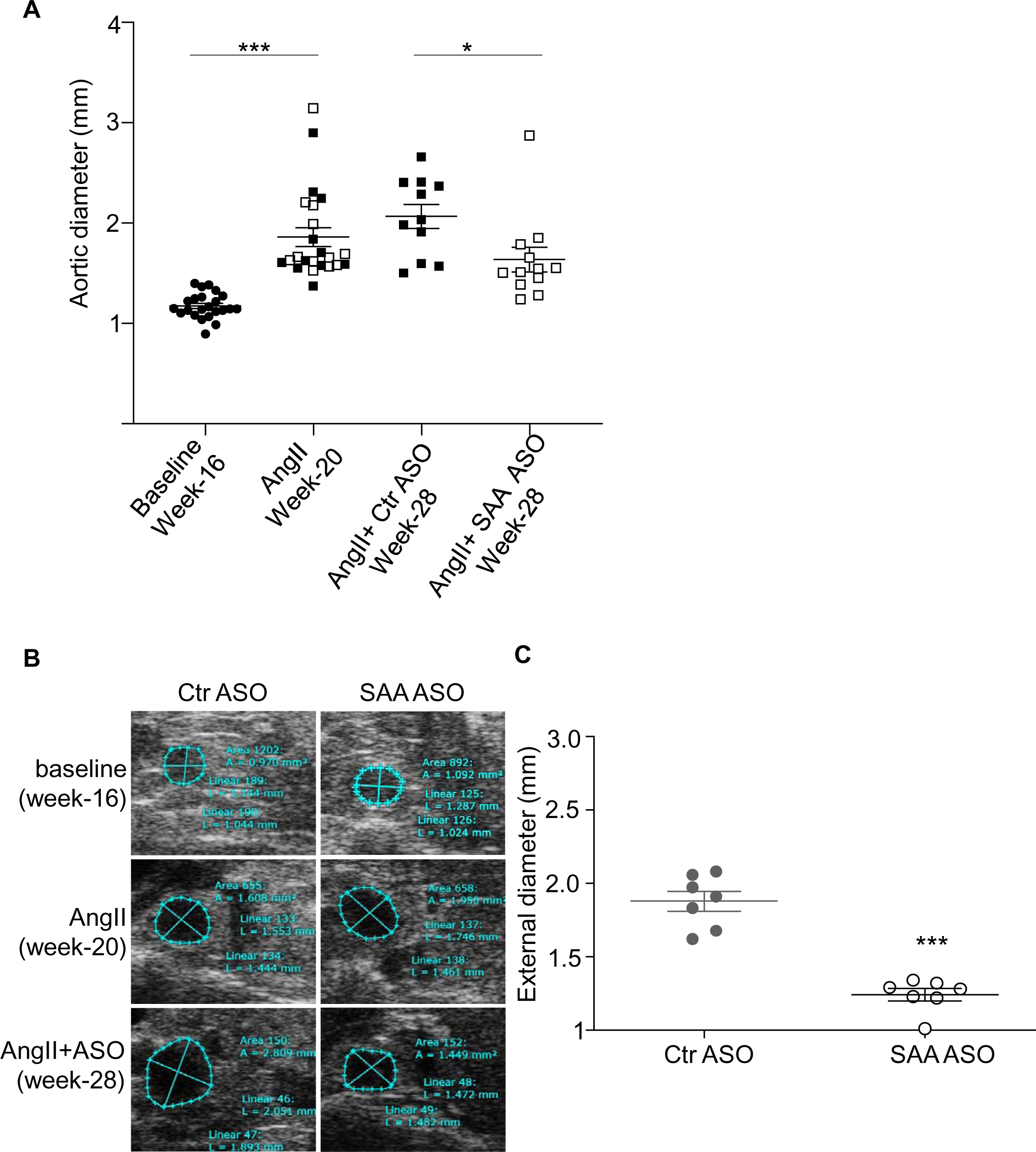
Mice treated with SAA ASO demonstrated a significant reduction in AAA progression compared to the mice treated with control ASO. (A) Abdominal aortas were assessed by in vivo ultrasound before (closed circle, week-16), after 4 weeks of AngII infusion (The maximal luminal diameters of the mice assigned to receive Ctr-ASO and SAA-ASO are shown by closed and open squares respectively, week-20) and after 12 weeks of AngII infusion along with a final 8 weeks of ASO treatment (Ctr-ASO-closed square and SAA-ASO-open square; week-28) to determine maximal luminal diameters in individual mice. (B) Representative US images of the aorta are shown. (C) the aortas were cleaned from a subset of animals in each group (aortas were chosen with maximal luminal diameters around the average in each group) for AAA assessment ex vivo by computer-assisted morphometric analysis to determine the maximal diameter of abdominal aortas. Each symbol represents one animal. *= P ≤ 0.05; and ***= P ≤ 0.001.

### Suppression of SAA is associated with decreased macrophage infiltration and matrix metalloproteinase (MMP) activity in abdominal aortas during AngII-induced AAA progression

Abdominal aortic tissue sections from mice treated with Ctr ASO or SAA ASO at the study end were examined for elastin breaks, macrophage infiltration, and MMP activity, three well-documented features of AAA ^20^. Elastin breaks were more prominent in the aortas of mice treated with Ctr ASO than in those treated with SAA ASO (representative images are shown in Fig. 5A). Aortic sections from Ctr ASO-treated mice demonstrated significantly more MMP activity than the sections from SAA ASO treated mice (Fig. 5A and Fig. 5B). Furthermore, macrophage infiltration, as evidenced by CD-68 immunostaining, was significantly greater in the aortic sections of mice treated with Ctr ASO than those of the SAA ASO treated mice (Fig. 5A and Fig. 5C). There was a significant reduction in SAA immunostaining in the aorta of mice treated with SAA ASO as compared to Ctr ASO treated mice (Fig. 5A and Fig. 5D). The specificity of in situ zymography experiments was confirmed by performing the staining procedure in the presence of an MMP inhibitor (Supplementary Fig. S1A) and the specificity of macrophage and SAA immunoreactivity was confirmed by staining sections in the absence of primary antibodies (Supplementary Fig. S1B). Thus, our data demonstrate that SAA promotes the progression of early AAA in mice, and suppressing SAA levels limits the progression of AAA and may even lead to improvement in aortic dilation in obese C57BL/6 mice.

**Figure 5.**
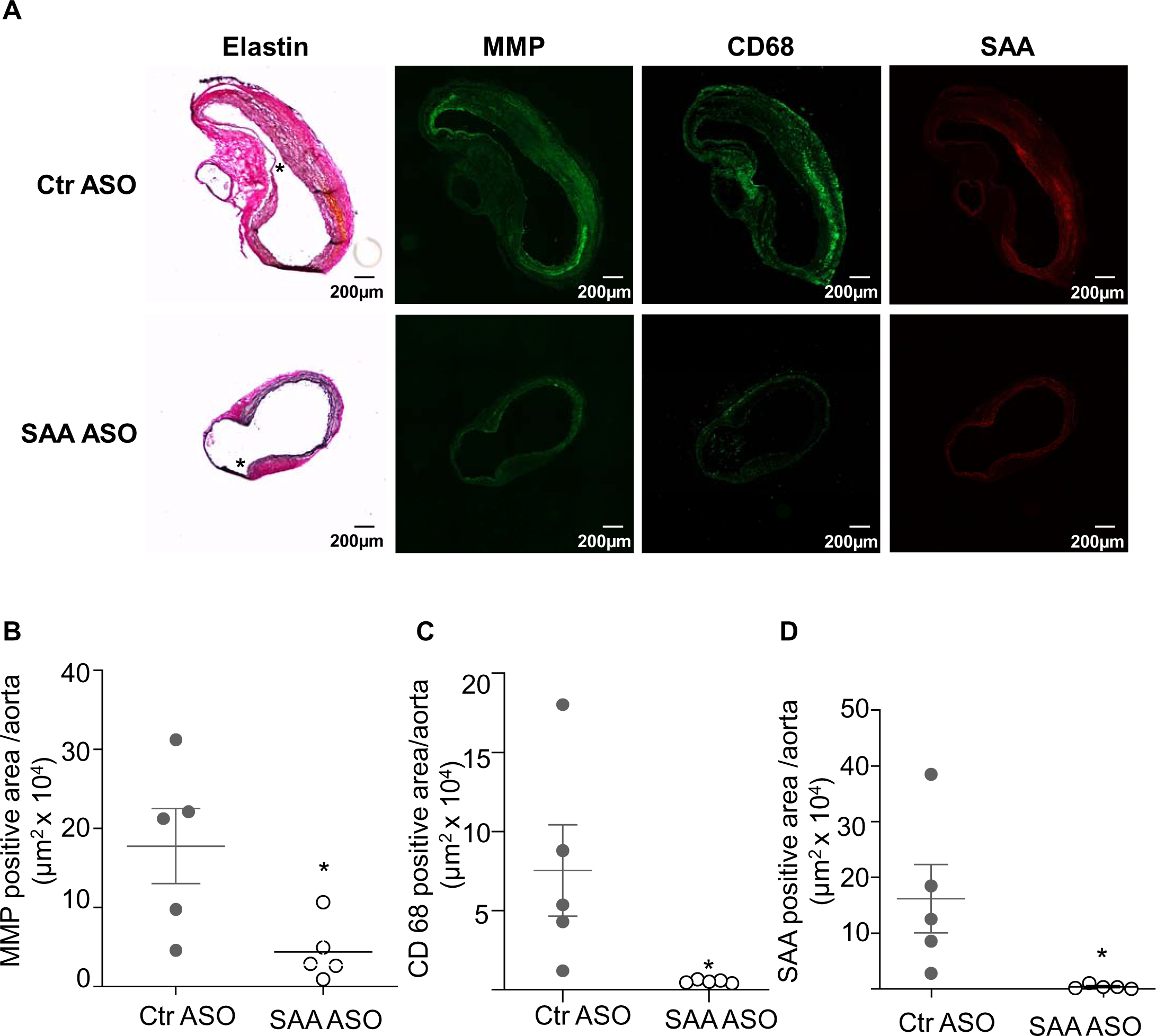
Mice treated with SAA ASO demonstrated a significant reduction in the matrix metalloproteinase (MMP) activity and accumulation of CD68-positive cells in the abdominal aortas. (A) Aortic sections of mice treated with AngII for a total of 12 weeks and with either control ASO (Ctr ASO) or SAA ASO for the final 8 weeks of AngII treatment were processed to detect elastin fibers (* denotes region with elastin breaks); MMP activity by in situ zymography (green fluorescence); macrophages with an anti-mouse CD68 antibody (green fluorescence middle panel), SAA in the aorta with an anti-mouse SAA antibody (red fluorescence) by immunostaining as described in methods, scale bar represents 200 μm. (B) MMP activity by in situ zymography was quantified from nine serial sections of abdominal aortas from individual mice, N=5 mice/treatment. Statistical analysis was performed using two-tailed t-tests, *P < 0.05. (C) CD68 immunostaining was quantified from nine serial sections of abdominal aortas from individual mice, N=5 mice/treatment. (D) SAA immunostaining was quantified from nine serial sections of abdominal aortas from individual mice, N=5 mice/treatment. Statistical analysis was performed using two-tailed t-tests, *P < 0.05.

**Figure 6.**
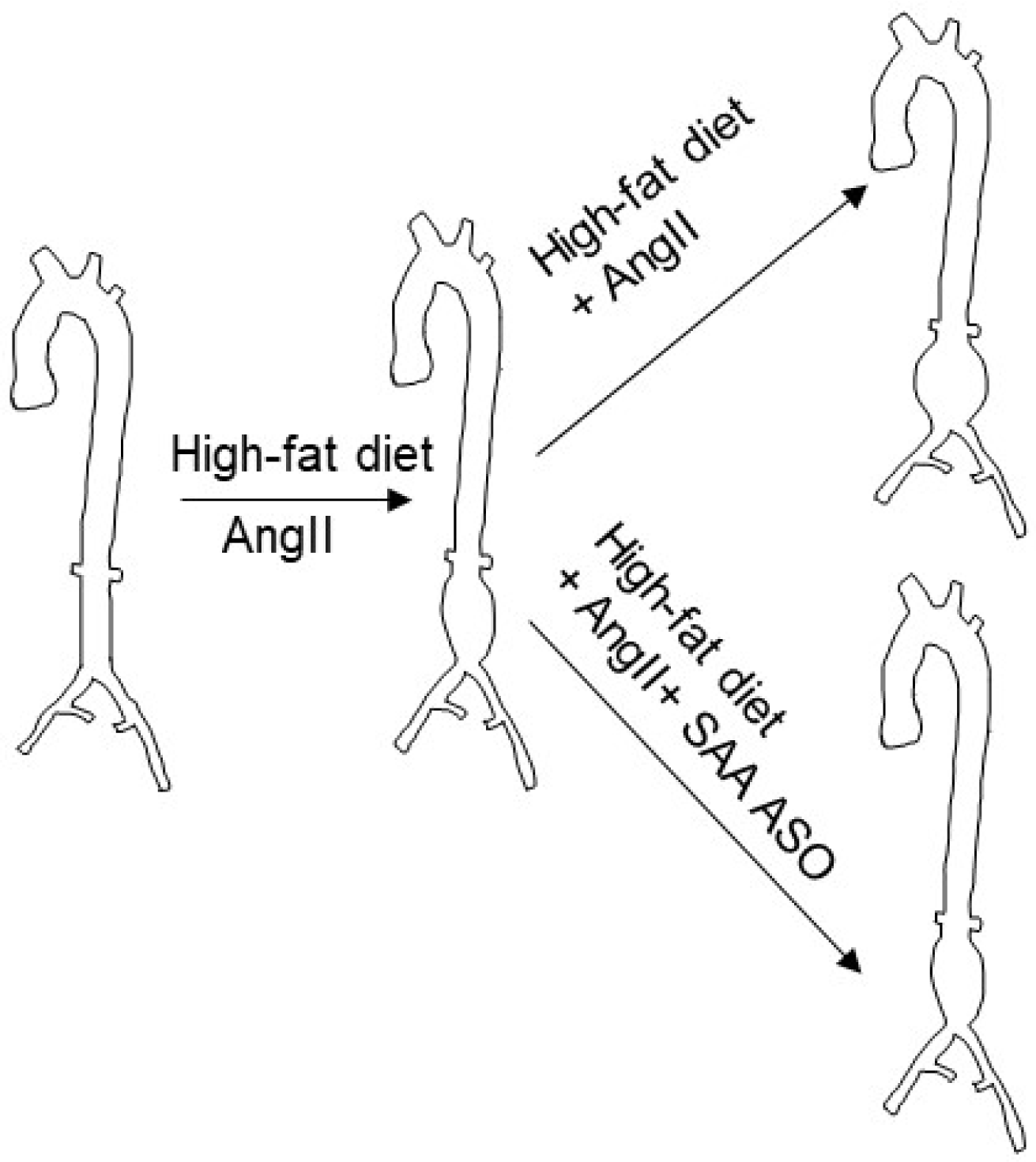
Suppression of SAA decreases the progression of AngII-induced AAA in obese mice. High-fat diet and AngII treatment increase SAA levels in circulation and induce the development of AAA in mice. Progression of early AAA in obese mice was significantly reduced when the mice were treated with ASO to suppress the expression of SAA compared to mice treated with control ASO.

## Discussion

AAA rupture is a prominent cause of death in adults, particularly males older than 65 years of age. The only available treatment for AAA is open or endovascular surgical repair. Though early detection and progression of the disease is possible by imaging and screening programs, surgical repair of small AAAs was not found to be beneficial ^21^, highlighting the urgent need for effective therapies that limit the progression of small AAAs. Our data demonstrate for the first time that targeting SAA may be beneficial for reducing the progression and future rupture of an early AAA.

The current study expands on results from our previous reports that SAA is essential for the development of AngII-induced AAA formation in both hyperlipidemic apoE^-/-15^ and obese C57BL/6 mice ^16^. In the current study, we investigated the protective effect of an antisense oligonucleotide inhibitor of SAA synthesis on the progression of early AngII-induced AAAs ^13^ in obese C57BL/6 mice. Obesity increases the risk for human AAAs ^22^ and enhances AngII-induced AAA formation in C57BL/6 mice ^14^. Both obesity and AngII treatment increase systemic SAA levels in mice ^16^. The liver is thought to be the primary source of circulating SAA in mice following AngII infusion. Although SAA expression is induced in adipose tissues following HFD feeding, its contribution to circulating SAA is negligible^16^. Thus, we opted to reduce plasma SAA levels in AngII-infused obese mice by using an N-acetylgalactosamine-conjugated ASO (GalNAc-ASO) that targets hepatic SAA to investigate whether systemic SAA plays a role in the progression of an already developed AAA. In this study, we considered mice with at least a 25% increase in the luminal diameter of the abdominal aorta to have an early AAA, and they were stratified into two groups to receive either control ASO or SAA ASO. Eight weeks of ASO treatment significantly decreased systemic SAA levels (Fig. 2A) and hepatic expression of SAA1.1 and SAA2.1 (Fig.2B and C). The ASO did not suppress SAA3 expression in the liver. SAA1.1 and SAA2.1 are the major acute-phase isoforms, and SAA3 is a minor isoform present in the circulation at significantly lesser levels than SAA1.1 and SAA2.1 ^10^. Our earlier studies indicated that the lack of SAA1.1 and SAA2.1 protects hyperlipidemic apolipoprotein E-deficient mice against AngII-induced AAA development ^15^. We have also demonstrated that deficiency of all three isoforms in obese mice abrogates the development of AngII-induced AAA in mice ^16^. However, whether SAA3 has, any independent role in AAA development or progression is unknown. Interestingly, SAA3 is not expressed in humans. Therefore, it is of limited interest from a clinical point of view. Consistent with our previous report ^16^, there were no significant differences in plasma cholesterol between the mice treated with control ASO and SAA ASO. In contrast to our earlier study, where there was a significant increase in mean systolic blood pressure in mice deficient in SAA1.1 and SAA2.1 compared to the wild-type mice ^15^, SAA ASO treatment did not cause a significant change in blood pressure. However, there was a trend for decreased blood pressure in the mice treated with SAA ASO compared to the control ASO group. The effect of SAA on blood pressure has not been thoroughly studied. Elevated blood pressure is a common feature in the well-established AAA model. However, AngII-induced AAA develops by mechanisms that are independent of blood pressure ^23^.

The major finding of the present study is that the administration of SAA ASO significantly reduced the progression of early AAA in mice (Fig.4 and 6). There was continued progression of AAA in mice between 4 weeks of AngII infusion to 12 weeks of AngII infusion in Ctr ASO-treated mice; however, the diameter decreased by 11% in the SAA ASO-treated mice (Fig.4A). Here, we show that suppression of systemic SAA is sufficient to arrest the progression of an early AAA. Deficiency of SAA did not cause any significant change in systemic inflammatory responses in AngII-induced AAA formation in both hyperlipidemic ^15^ and obese ^16^ mouse models; in both studies, alterations in the severity of AAA were associated with alterations in the amount of SAA detected in aneurysmal tissue. Thus, SAA derived from either source, the liver or adipose tissue, may deposit in the vessel wall and have a pathological effect.

Our data provide some insights into the mechanisms by which SAA is involved in the progression of AAA. We have previously demonstrated that SAA increases MMP activity in the aorta ^15^; MMP activities, macrophages, and elastin breaks appeared absent in the abdominal aortas of AngII-infused SAA-deficient mice^16^. MMP-mediated extracellular matrix destruction is considered one of the major causes of the development and progression of AAA^24, 25^. MMP inhibition has emerged as a potential pharmacotherapeutic approach to limit the development and progression of AAAs^26^. However, most synthetic MMP inhibitors failed *in vivo,* likely because of low bioavailability and apparent adverse events^27^. Our data indicated that SAA ASO treatment significantly suppressed MMP activity in the aorta of mice with AAA, paralleling a significant reduction in CD68-positive macrophages in the aorta (Fig. 5A, B, and C). Thus, consistent with our previous study on AAA development in mice^15, 16^, the current study indicated that SAA plays a role in enhancing MMP activities in the aorta during AAA development and progression, leading to increased macrophage infiltration and elastin breaks.

In summary, we demonstrated that the administration of SAA ASO suppresses systemic SAA levels in mice and limits the progression of early AAA in mice, possibly by decreasing MMP activities in the affected aortic regions. Our study shows that inhibition of SAA may be an effective therapeutic strategy for AAA progression.

## Conflict of interest

The authors declare that they have no conflicts of interest with the contents of this article.

## Financial support

This work was supported by Department of Veterans Affairs BX004275 (to LT and DK) and National Institutes of Health Grant HL147381 (to LT and PS).

## Author Contributions

Conceptualization: Nancy R. Webb, Lisa R. Tannock, Preetha Shridas.

Methodology: Ailing Ji, Nancy R. Webb, Lisa R. Tannock, Preetha Shridas.

Validation: Ailing Ji, Preetha Shridas.

Formal analysis: Ailing Ji, Preetha Shridas.

Investigation: Ailing Ji, Andrea C. Trumbauer, Victoria P. Noffsinger, Luke W. Meredith, Preetha Shridas.

Resources: Adam E. Mullick, Frederick C. De Beer, Dennis G Karounos.

Data curation: Ailing Ji, Preetha Shridas.

Writing – original draft: Ailing Ji, Preetha Shridas.

Writing – review & editing: Ailing Ji, Nancy R. Webb, Lisa R. Tannock, Preetha Shridas.

Visualization: Ailing Ji, Preetha Shridas.

Supervision: Preetha Shridas, Lisa R. Tannock.

Project administration: Preetha Shridas, Lisa R. Tannock.

Funding acquisition: Dennis G Karounos, Preetha Shridas, Lisa R. Tannock

## Acknowledgments

The studies were supported with resources and facilities provided by the Centers of Biomedical Research Excellence (COBRE) at the University of Kentucky, which was supported by an Institutional Development Award (IDeA) from the National Institute of General Medical Sciences of the National Institutes of Health under grant number P30 GM127211.

## Notes

### Competing Interest Statement

The authors have declared no competing interest.

### Summary of Updates

Presented Figure 4 A in a different format. Added a graphical abstract as Figure 6 and added a supplemental figure.

