## Supplemental Figure for "Antisense Oligonucleotide Targeting Hepatic Serum Amyloid A Limits the Progression of Angiotensin II-Induced Abdominal Aortic Aneurysm Formation": Shridas et al Spplementary data.pptx

### Slide 1
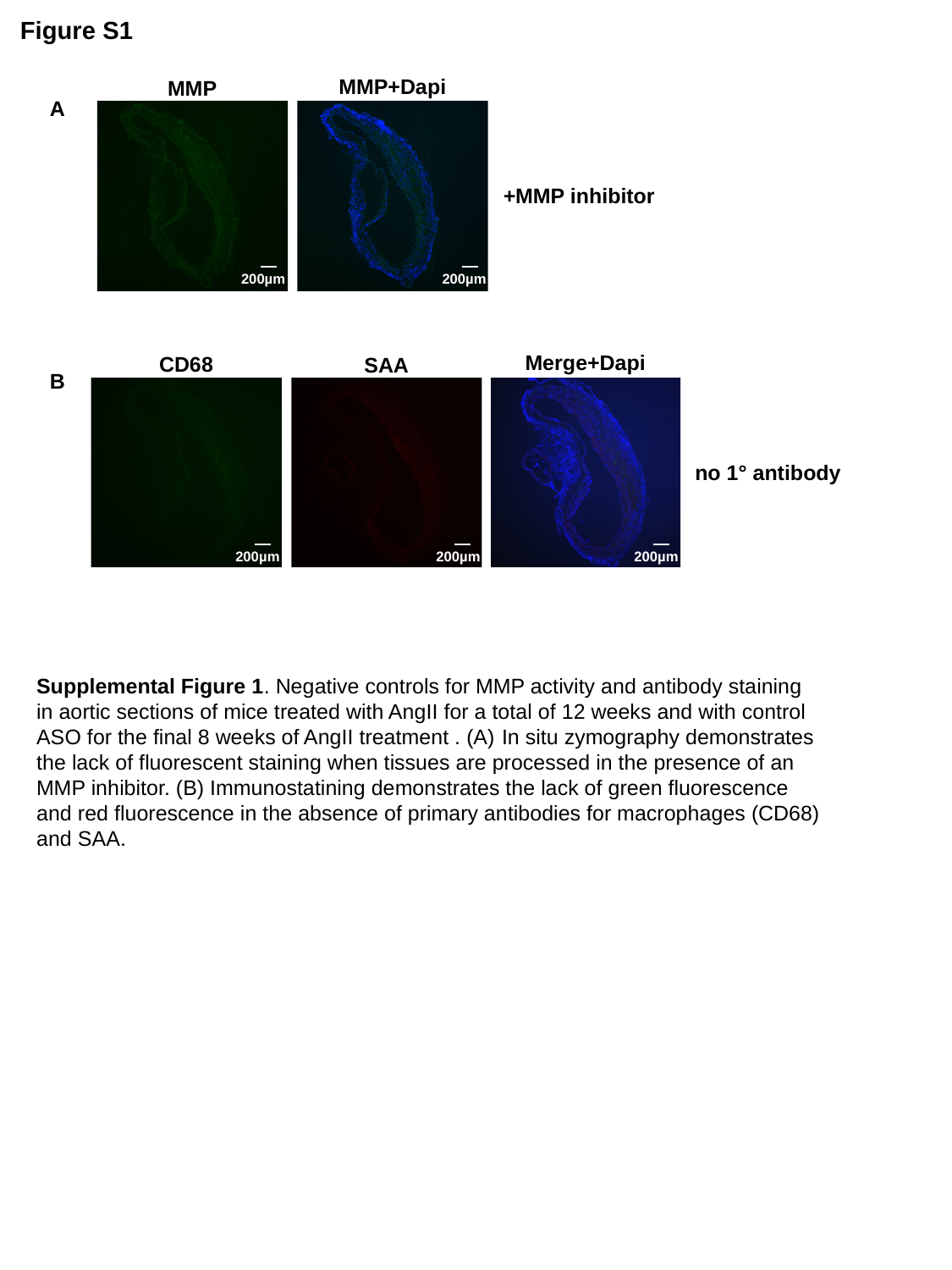

Figure S1
MMP+Dapi
MMP
A
+MMP inhibitor
200µm
200µm
Merge+Dapi
CD68
SAA
B
no 1° antibody
200µm
200µm
200µm
Supplemental Figure 1. Negative controls for MMP activity and antibody staining in aortic sections of mice treated with AngII for a total of 12 weeks and with control ASO for the final 8 weeks of AngII treatment . (A) In situ zymography demonstrates the lack of fluorescent staining when tissues are processed in the presence of an MMP inhibitor. (B) Immunostatining demonstrates the lack of green fluorescence and red fluorescence in the absence of primary antibodies for macrophages (CD68) and SAA.
